# Single Cell Analysis of pBMCs of Psoriasis patients reveals distinct CD4+ T cell phenotypes associated with response to IL-23 blockade

**DOI:** 10.1101/2025.11.04.686122

**Authors:** Kathryne E. Marks, Anina Sun, Ifeoluwakiisi Adejoorin, Lourdes Perez Chada, Anna Perillo, Leanne Barrett, Michael Garshick, James Krueger, Joseph F. Merola, Marcelo DiCarli, Deepak A. Rao, Brittany N. Weber

**Affiliations:** Division of Rheumatology, Inflammation, Immunity, Brigham and Women’s Hospital and Harvard Medical School, Boston, MA, USA; Division of Cardiovascular Medicine, Brigham and Women’s Hospital, Harvard Medical School, Boston, MA, USA; Leon H. Charney Division of Cardiology, New York University Grossman School of Medicine, New York, New York; Laboratory for Investigative Dermatology, The Rockefeller University, New York, NY, USA; Department of Dermatology and Department of Medicine, Division of Rheumatology, UT Southwestern Medical Center and O’Donnell School of Public Health, Dallas, TX, USA; Cardio-Rheumatology and Cardio-Dermatology Program, Division of Cardiology, Department of Internal Medicine and Dermatology, University of Texas Southwestern Medical Center, Dallas, Texas, USA

## Abstract

The use of IL-23 inhibitors (IL-23i) for psoriatic diseases has resulted in significant improvement in disease symptoms for many patients. The extent of disease improvement following IL-23 blockade varies across patients with psoriasis; however, the immunologic factors associated with a good or poor response to IL-23 blockade remain unclear. Here, we utilized peripheral blood mononuclear cells (pBMCs) collected through the MINIMA clinical trial (NCT04271540) and applied single-cell RNA sequencing to profile circulating immune populations from 27 patients with psoriasis or psoriatic arthritis, aiming to identify cellular features associated with a response to IL-23 blockade. We identified populations of CD4^+^ T cells whose abundance in circulation before treatment was associated with improved skin disease following IL-23i treatment. Circulating CD4^+^ T cells that demonstrate transcriptomic features of Th1-like cells were associated with better psoriasis skin improvement and had transcriptomic signatures resembling T cells identified in psoriasis lesional skin. Further, decrease in levels of IL-17A and IFNγ in serum each correlated with improvement in PASI levels. These results suggest that immune cell features detectable in blood may be informative in identifying patients with psoriasis who are likely to have a robust clinical response to IL-23 blockade.

## Introduction

Psoriasis is a common autoimmune disease characterized by red, raised scaly lesions on skin (1, 2). Patients with psoriasis experience systemic inflammation that can predispose to the development of psoriatic arthritis and accelerated development of cardiovascular disease (1-3). There are several treatments available for psoriasis, including inhibitors of TNF; however, inhibitors of IL-17 or IL-12/23 have been particularly effective for skin disease (2, 4). The regular use of dedicated IL-23 inhibitors (IL-23i) is relatively recent, and their effect in modulating the systemic immune system is not yet clear.

IL-17 and IL-23 inhibitors inhibit the IL-23/Type 17 T cell axis by inhibiting the differentiation or function of pathogenic Th17 cells (2, 5). Th17 cells vary in their ‘pathogenic’ potential depending in part on additional factors they produce, with co-production of IFN-γ associated with pathogenic cells and co-production of IL-10 associated non-pathogenic cells. Non-pathogenic or homeostatic Th17 cells are important for mucosal homeostasis and defense against fungal infections, while pathogenic Th17 cells are more strongly associated with inflammatory disease (6, 7). Pathogenic Th17 cells, especially those that co-express IFNγ and IL-23R, have been implicated in several autoimmune diseases, including psoriasis, multiple sclerosis, and ulcerative colitis (8-10).

A range of IL-17-producing T cells accumulate within psoriasis skin lesions (11, 12); yet, recent findings have shown IL-23 blockade with rizankizumab strongly and preferentially reduces the abundance of IL-17A^+^IFNγ^+^ and IL17F^+^IL10^-^ Th17 cells in skin. In contrast, the proportion of IL17A^+^IL17F^+^ cells, which have low expression of IL-23R, increases following IL-23 blockade (12). Whether the systemic balance of pathogenic to non-pathogenic Th17 cells is associated with response to IL-23 blockade is unclear.

We aimed to identify T cell populations that are associated with clinical response to tildrakizumab, an FDA approved inhibitor of the p19 subunit of IL-23, in patients with active psoriasis. To do this, we utilized peripheral blood mononuclear cells that were collected through a mechanistic clinical trial (MiNIMA, NCT04271540). In this study, patients with moderate-severe psoriasis and elevated cardiovascular risk not currently on any psoriatic treatment were enrolled and assessed at baseline and after 24 weeks of therapy. By phenotyping subsets of circulating CD4+ T cell populations with single cell RNA sequencing, we show that higher proportions IFNγ-Th17 cells in blood are associated with a weaker clinical response to tildrakizumab. We further compare the response-associated T cells with transcriptomes of Type 17 T cells in psoriasis lesional skin to demonstrate their similarity to skin T cells.

## Results

### Patient characteristics and study setup

Patients with moderate to severe psoriasis were recruited to the MIcrovascular dysfuNction In Moderate-severe Psoriasis (MINIMA) clinical trial at Brigham and Women’s Hospital (BWH)[NCT04271540]. Briefly, MINIMA was a clinical trial designed to determine if Tildrakizumab improves coronary vascular function and coronary flow reserve, as measured by noninvasive imaging with cardiac positron emission tomography after 6 months of treatment with Tildrakizumab compared to baseline. If the patient was on a psoriasis therapy, they underwent a required washout period based on the half-life of the prior therapy before starting tildrakizumab for psoriasis treatment. Skin severity was measured throughout the trial using the psoriasis area and severity index (PASI), as well as physician global assessment (PGA) on a scale of 0-5 with 5 being the most severe assessment by board-certified dermatologists (13).

PBMCs were collected before and 24 weeks after treatment initiation with tildrakizumab (Figure 1A). We included an additional two patients who were enrolled in a similar yet more pragmatic clinical study performed during the same time period that examined patients prior to and following 24 weeks of therapy with biologics that similarly target IL-23 inhibition. The two patients included from this study had samples collected before and after treatment with other similar IL-23 inhibitors, risankizumab and guselkumab. A summary of the patients is in Table 1. Patients who had not previously been treated with an IL-23 or IL-17 inhibitor or were required to have a wash-out period as above.

**Figure 1.**
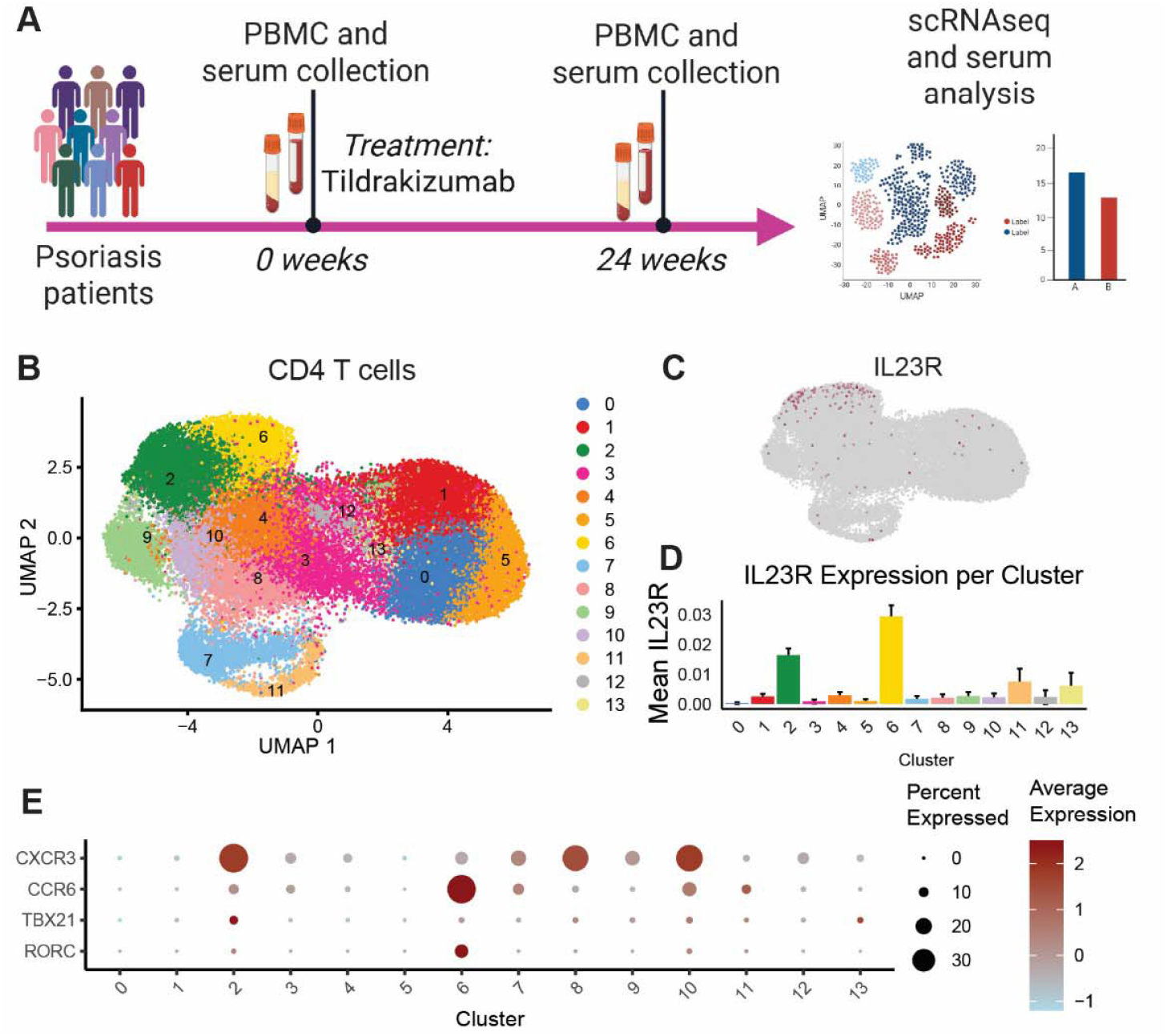
Identification IL-23R^+^ CD4 T cells in blood of patients with psoriasis. A) Schematic of the MINIMA and ACTION study designs. B) UMAP of scRNAseq of CD4 T cells from blood. C) Expression of IL23R within CD4^+^ T cells. D) Mean IL23R expression within CD4^+^ T cells. E) Expression of indicated genes within CD4^+^ T cells.

**Table 1.** scRNAseq samples.

| Table 1. scRNAseq samples |  |
| --- | --- |
| Pre/Post paired samples | 27 |
| FollowUp | ~24 weeks |
| Age median (range) | 63 (31-79) |
| Female median (%) | 10 (37%) |
| PsA (%) | 6 (22%) |
| Treated with tildrakizumab | 25 |
| Treated with risankizumab | 1 |
| Treated with guselkumab | 1 |
| DeltaPASI median (range) | -7.8 (-0.9--17.7) |
| PASI improved by 75% | 13 (48%) |

Among the cohort, the median age was 63 (IQR), 37% female, and 6 patients were diagnosed with psoriatic arthritis. All patients’ skin symptoms improved to some extent with a median change in PASI of -7.8 (IQR -0.9 - -17.7, p < 0.000001) (Table 1).

### scRNAsequencing of blood from psoriasis patients before and after tildrakizumab treatment

We performed single cell RNA sequencing (scRNA-seq) on total live PBMCs according to previously described methods (14, 15), followed by quality control and batch correction using Harmony (14). Following QC, 130,000 cells from PBMCs from 27 unique patients were available for analysis, including cells from paired pre- and post-treatment timepoints from 26 patients.

Cells from PBMCs clustered into the expected major immune cell lineages (Figure S1A). The proportions of naïve and memory CD4^+^ T cells, naïve and memory CD8^+^ T cells, NK cells, B cells, and myeloid cells were overall stable comparing pre-treatment and post-treatment timepoints (Figure S1B).

### Association of circulating T cells with clinical response to tildrakizumab

We focused analyses on CD4^+^ T cells, which contain cells that are sensitive to IL-23 (2, 16) (Figure 1B). In this dataset we observed two clusters of CD4^+^ T cells enriched in IL23R expression (clusters 2,6) (Figure 1C,D). These clusters contained cells with phenotypes reflecting Th1 and Th17 CD4^+^ T cell subsets, with expression of TBX21 and CXCR3, and RORC and CCR6, respectively (Figure 1E).

The IL23R^+^ clusters 2 and 6 were selected for further analysis. Subclustering the cells within these clusters resulted in 5 subclusters of cells with varying levels of Th17 and Th1 phenotypes (Figure 2A). To test whether any of these IL23R+ cell populations were associated with clinical response to IL-23i, we used covarying neighborhood analysis (CNA) to identify fine grained cell neighborhoods associated with disease activity (17). CNA analysis testing the association of IL-23R enriched cell phenotypes at the pre-treatment timepoint with PASI levels at the post-treatment timepoint identified cells significantly associated with higher or lower disease activity after treatment (Figure 2B). Cells on the right side of the UMAP, mostly within cluster 0, were significantly associated with higher PASI after treatment, while cells on the left side of the UMAP, mostly within clusters 1 and 2, were associated with lower PASI after treatment. Cluster 0 contained cells expressing high levels of RORC, CCR6, CCL20, and low levels of CXCR3, while clusters 1 and 2 contained higher expression of TBX21, IFNγ, and CXCR3 (Figure 2C, S2A).

**Figure 2.**
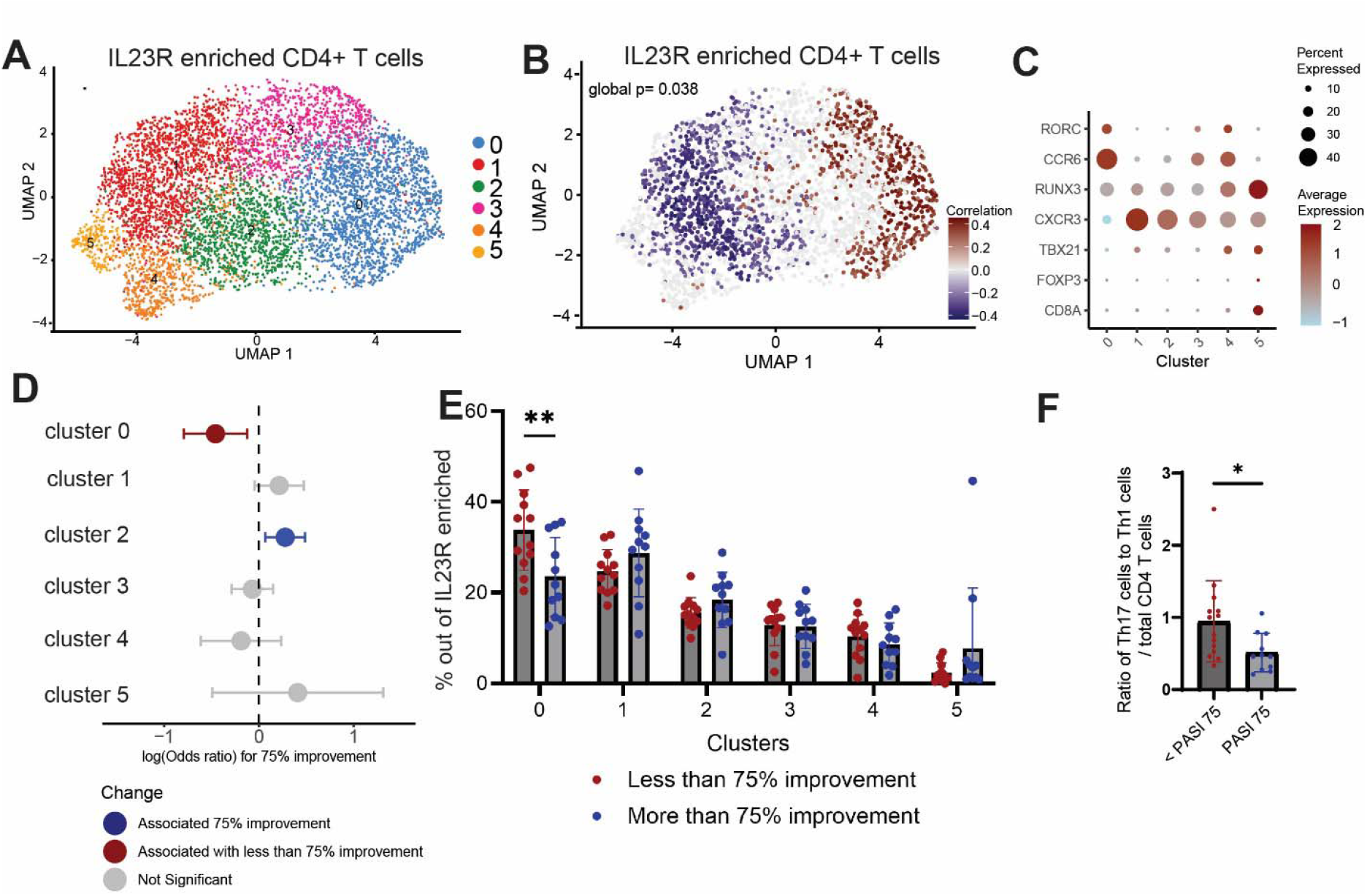
Association of CD4^+^ T cell subsets at baseline with extent of improvement in PASI. A) UMAP of IL23R enriched CD4^+^ T cells B) Covarying Neighborhood Analysis of IL23R enriched CD4^+^ T cells before treatment for association with PASI after treatment. C) Gene expression in IL23R enriched CD4^+^ T cells D) MASC analysis for association with PASI 75 among IL23R enriched cells before treatment. Analysis includes covariates of age and sex. E) Proportion of each cluster divided by PASI 75. F) Ratio of Th17 cells (cluster 6) to Th1 cells (cluster 2) out of total CD4^+^ T cells.

In a complementary analysis using a dichotomous definition of treatment response, we tested for IL23R+ cell populations associated with attainment of a PASI 75, a frequently used benchmark of treatment success (18). We implemented mixed-effects modeling of associations of single cells (MASC), which identifies cell clusters associated with a clinical variable using a mixed-effect logistic regression model at a single-cell level (19). Cluster 0 was significantly associated with failure to achieve PASI 75, while cluster 2 was significantly associated with achieving a PASI 75 (Figure 2D). Cluster 1, which was transcriptomically similar to cluster 2, also showed a trend towards association with achievement of PASI 75. Consistent with these results, the frequency of cluster 0 out of all IL-23R enriched cells was significantly higher in patients who reached PASI 75 compared to patients who did not (Figure 2E). In addition, we also dichotomized patients by those who had a decrease in physician global assessment (PGA) of 3 or greater. Consistent with results using PASI 75, cluster 0 was more highly represented in patients with a PGA reduction of less than 3 (Figure S2B). We then compared the ratio of cells in the Th17 cluster to cells in the Th1 cluster among total CD4^+^ T cells. Patients who did not achieve PASI 75 after IL-23i had a higher ratio of Th17 to Th1 cells compared to patients who reached PASI 75 (Figure 2F).

A similar analysis of cells from the post-treatment samples showed a similar trend; however, the response level association was no longer significant (Figure S2C). Analysis of cluster proportions in pre-vs post-treatment samples also trended towards a decrease in clusters 1 and 2 and an increase in cluster 0, but varied between patients and was not statistically significant (Figure S2D).

### Characterization of response associated cells

To further characterize the transcriptomic identities of the cell populations associated with clinical improvement level after IL-23i treatment, we compared the IL-23R enriched cell transcriptomes to T cells that are present in psoriasis skin lesions using previously reported data (11). We created gene scores from genes highly expressed by IL-17-producing lesional skin T cells, including IL-17A^+^, IL-17A^+^IFNγ^+^, IL-17A^+^IL17F^+^, IL-17F^+^, IL-17F^+^IL-10^+^ T cells (Supplementary Table 2). Cluster 0, the IL-23R enriched cell cluster associated with a weak response to IL-23i, had low expression of gene scores for IL-17A^+^ and IL-17A^+^IFNγ^+^ T cells and higher expression of gene scores for IL-17A^+^IL17F^+^ and IL-17F^+^ T cells (Figure 3A). Importantly, a previous study demonstrated that skin T cells of the IL-17A^+^IL-17F^+^ phenotype are retained following IL-23i, while the IL-17A^+^IFNγ^+^ T cells are dramatically reduced (12).

**Figure 3.**
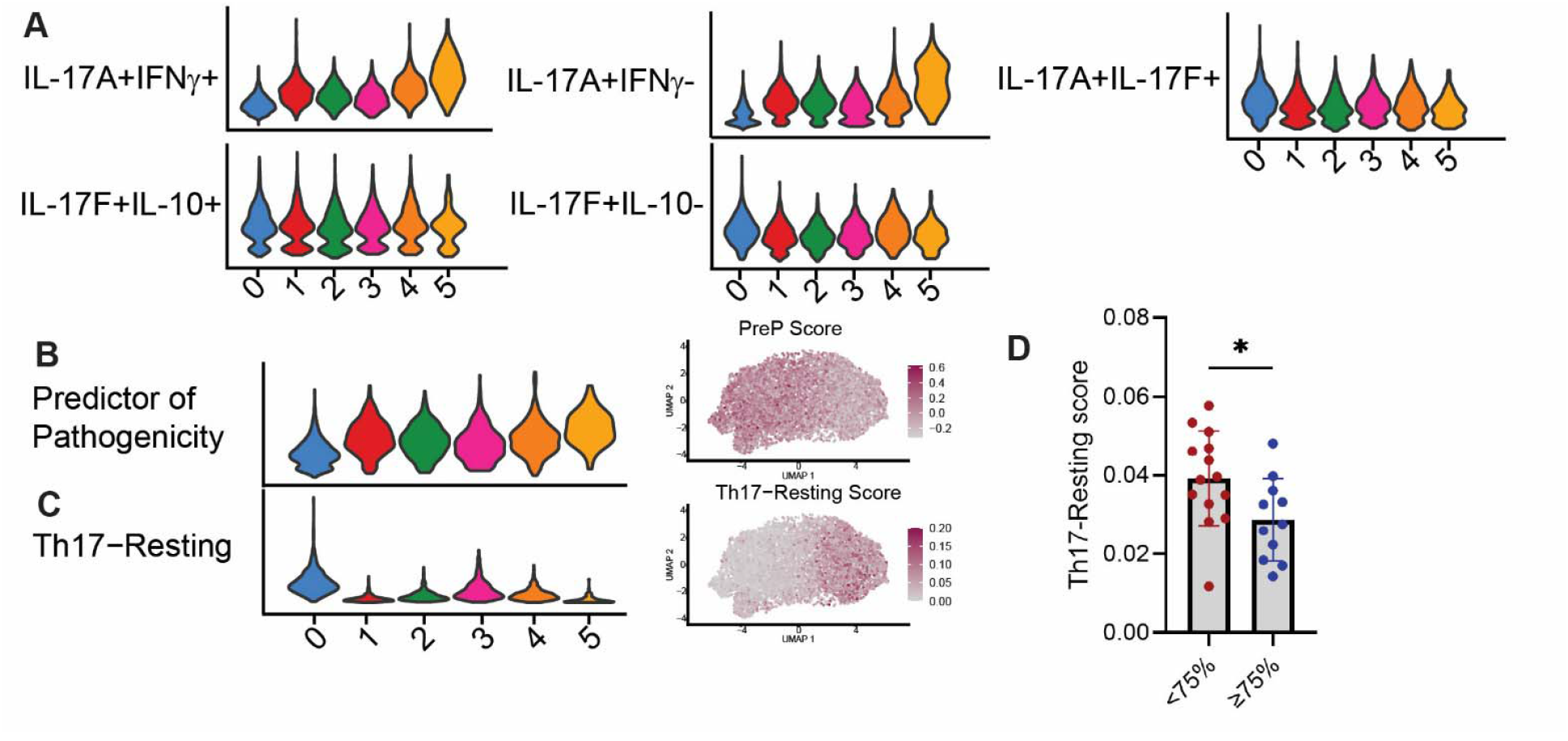
Comparisons of transcriptomes of IL23R enriched cells to IL-17 producing cells in psoriasis skin. A) Gene scores (Supplementary Table 2) applied to the IL23R enriched CD4+ T cell clusters. B) PreP score (Supplementary Table 3) applied to IL23R enriched CD4+ T cell clusters. C) Th17-Resting TCAT score applied to IL23R enriched CD4 T cells. D) Th17-Resting score divided by PASI 75. ^*^p < 0.05 by Mann Whitney test.

These results suggested that cluster 0 most aligns with a non-pathogenic, homeostatic Th17 cell population. Consistent with this idea, expression of a previously reported ‘predictor of pathogenicity’ gene signature was low in cluster 0 and higher in clusters 1 and 2 (Figure 3B; Supplementary Table 3) (9).

We further characterized the IL-23R enriched clusters by using TCAT, a recently developed algorithm to define characteristic T cell phenotypes and states (20). Cluster 0 was the only cluster that was characterized by expression of the genes belonging to the Th17-Resting phenotype (Figure 3C). Due to this distinctive set of features of cluster 0, we asked whether the Th17-Resting score itself was associated with clinical response to IL-23i therapy. Indeed, the Th17-resting score was significantly higher in pre-treatment blood IL-23R enriched cells from patients who did not achieve a PASI 75 as compared to patients who achieved a PASI 75 (Figure 3D).

### Serum levels of IL-17 and IFNγ and response to tildrakizumab treatment

To further characterize the correlates of treatment response in blood, we analyzed serum from patients before beginning tildrakizumab treatment. Comparison of serum levels of IL-17A and IFNγ with proportion of cells in cluster 0 or 2 of IL23R enriched CD4^+^ T cells revealed that pre-treatment levels of IFNγ were correlated with the proportion of cells in cluster 2. IL-17A levels, however, were not significantly correlated with any cluster including cluster, possibly implying IL-17A is produced by additional cells than those in cluster 0 (Figure 4A-D). Overall, pre-treatment levels of IL-17A in serum were correlated with disease activity by PASI, while pre-treatment levels of IFNγ were not (Figure 4E-F). Both the change in IL-17A and IFNγ levels in serum were correlated with the decrease in PASI (Figure 4G-H). Comparisons of serum levels before and after treatment revealed that IFNγ slightly, but significantly decreased from IL-23i, while IL-17A only trended towards a decrease from treatment.

**Figure 4.**
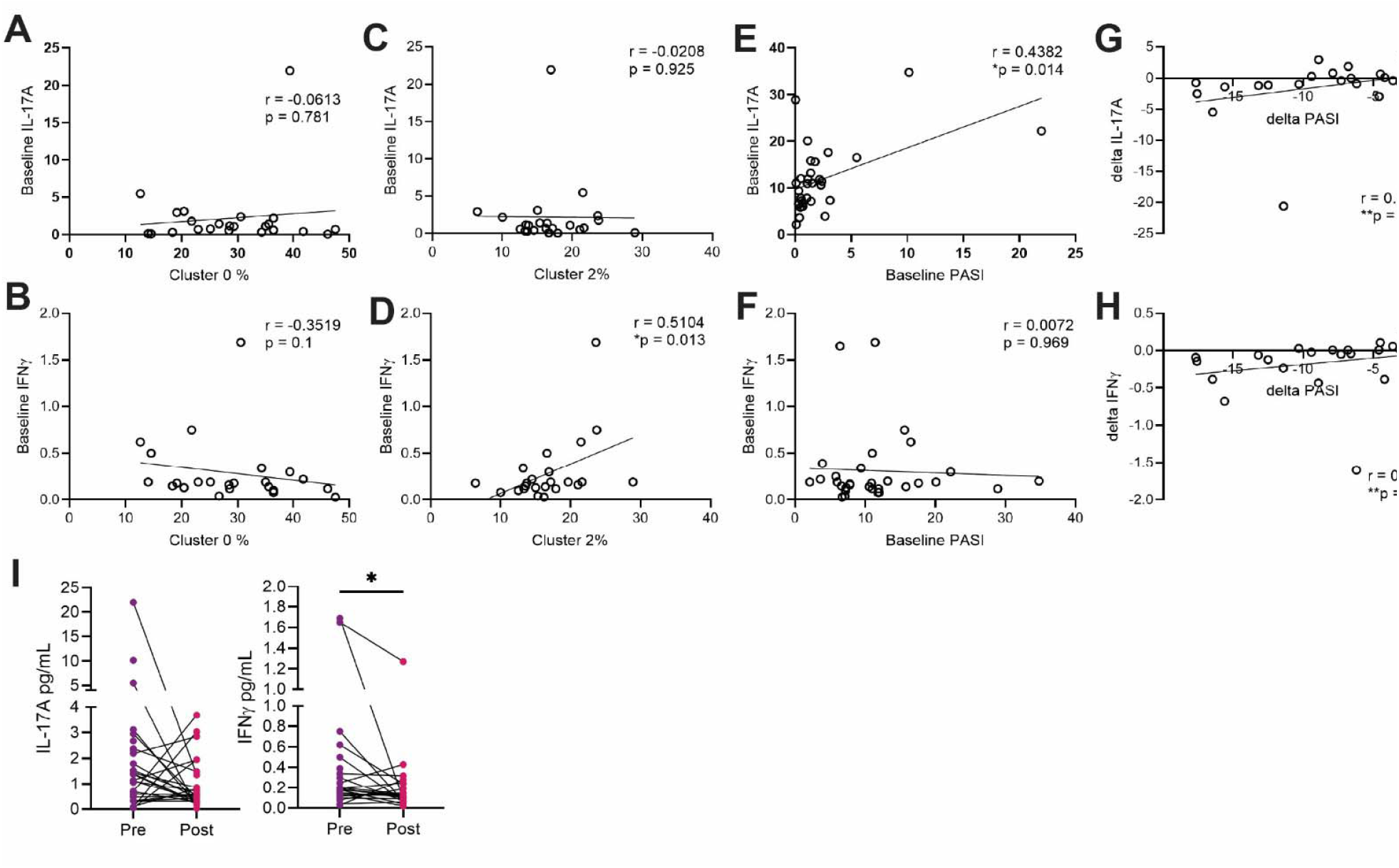
Associations of levels of IL-17A and IFNg at baseline with PASI. A) Spearman correlation of baseline IL-17A B) or IFNg with cluster 0 in IL23R enriched CD4+ T cells, C) Baseline IL-17A and D) IFNg with cluster 2 in IL23R enriched CD4+ T cells, D) Baseline IL-17A or E) IFNg with baseline PASI with baseline PASI, F) change in IL-17A and F) change in IFNg with change in PASI. I) Overall changes in IL-17A and IFNg pre vs post. ^*^p<0.05 by Wilcoxon matched pairs test.

## Discussion

In this study, we utilized scRNA sequencing to assess cellular phenotypes associated with clinical response in psoriasis patients treated with IL-23 blockade. We identified that proportion of homeostatic, or “resting”, Th17 cells at baseline are negatively associated with likelihood of a good response to tildrakizumab therapy for psoriasis. Conversely, circulating T cells with Th1-like phenotypes are positively associated with good response to IL-23 blockade.

By comparing transcriptomic signatures of circulating T cell subsets to those in psoriasis skin, we demonstrated that the population of circulating Th17 cells that was negatively associated with clinical response expressed low levels of a gene signature representative of pathogenic IL17A^+^ IFNγ^+^ cells, which have been demonstrated to be effectively reduced in skin by IL-23 blockade. Thus, we hypothesize that a lower proportion of CD4^+^ T cells that are affected by IL-23 blockade may indicate a lower likelihood of a good response to IL-23 blockade. Detection of these T cells in circulation may provide an accessible method to quantify the extent of IL-23-driven T cell phenotypes systemically in patients with psoriasis.

Our longitudinal scRNA-seq analyses of circulating T cells evaluated before and after IL-23 blockade did not identify consistent shifts in T cell subset proportions or transcriptomic signatures, even though both clinical disease and circulating IFNγ levels were reduced following IL-23 blockade in this cohort. It is possible that concurrent changes in T cell differentiation, migration into tissues, and exit from tissues make it challenging to detect consistent changes longitudinally in blood. Future studies should determine whether or not there are observable systemic shifts in circulating immune cells in other immune mediated inflammatory diseases for which IL-23 blockade is effective such as ulcerative colitis (21).

The patients included in our analysis were part of a clinical trial that included a treatment wash-out period prior to treatment with IL-23 blockade. Our study, however, is limited by the number of patients in the study and by a single method of analysis using scRNAseq. Additionally, we measured clinical response by PASI or global assessment at 6 months post initiation of IL-23 blockade. This timepoint may have missed further improvement in PASI after the primary endpoint timepoint. The response rate to tildrakizumab in this study is lower than in other reported studies of IL-23 blockade (22, 23). The intermediate response rate provided the ability to compare groups with and without a PASI 75 response with adequate statistical power; however, the applicability of this signature to other IL-23 blocking agents will require further study.

In summary, the use of scRNA-seq profiling of immune cells in blood of patients with psoriasis in the setting of a clinical trial enabled interrogation of fine-grained phenotypes of CD4^+^ T cells and their relationship to clinical response to a biologic therapy. These analyses identified a link between baseline levels of circulating Th17 cell subsets and psoriasis response to IL-23i treatment. Further study will clarify the optimal way to easily identify this phenotype in blood with the goal of advancing personalized treatment for psoriasis.

## Methods

### Study approval and design

MiNIMA is an open-label, single-center, single arm pilot study (NCT04271540). Patients with moderate-to-severe psoriasis and evidence of cardiovascular risk factors were recruited from Skin and Related Musculoskeletal Diseases Clinic and the Rheumatology Clinic from the Brigham and Women’s Hospital. ACTION was a pragmatic single-center study at Brigham and Women’s Hospital that recruited patients with moderate-to-severe psoriasis and evidence of cardiovascular risk factors who were starting an IL-23 or IL-17 inhibitor on clinical grounds by the prescribing physician for psoriasis treatment. After giving written informed consent to participate in the study and fulfilling criteria for inclusion, patients were scheduled for baseline cardiac PET/CT, and dermatology visit. Labs were drawn at the baseline visit and at the 24-week final cardiovascular visit.

Human subjects research was performed in accordance with the Institutional Review Board at Mass General Brigham. Blood samples were obtained as part of the MIMIMA or ACTION trials from patients with moderate to severe psoriasis before and 24 weeks after receiving tildrakizumab or risankizumab or guselkumab in the case of ACTION. Demographics of each patient can be found in Supplementary Table 1. Mononuclear cells from PBMCs were isolated using SepMate (STEMCELL technologies) tubes and cryopreserved in 10% DMSO in FBS and stored in liquid nitrogen. Serum was collected and stored in -80C.

### Single cell RNA sequencing

Cryopreserved PBMCs were thawed, washed, counted, fc blocked with FcX (Biolegend), and labeled with hashtag antibodies (Biolegend) in batches of 14 samples each. Baseline and post treatment samples were sequenced in the same batch. Propidium iodide negative cells were sorted by a BD Aria sorter. Cells were then stained with TotalSeq-C Human Universal Cocktail (Biolegend) according to manufacturer’s protocol. The cells were then encapsulated and the library was prepared at the BWH Center for Cellular Profiling via the 10X Genomics pipeline, with ∼72K cells loaded per run with 85-98% viability.

### Single cell RNA sequencing analysis

Samples were demultiplexed using hashtags antibodies before initial quality control which included removal of doublets and cells containing more than 15% mitochondrial content. Analysis was completed in the Seurat R package (V4)(24). Batch correction was completed using Harmony (14) for both batch and individual patient.

Further analyses included gene expression analysis of genes using Seurat AverageExpression, DotPlot, and FeaturePlot functions. Mixed-effects modeling of associations of single cells (MASC)(19) and Covarying Neighborhood Analysis (CNA)(17) were applied to IL23R enriched CD4+ clusters as previously described. The Seurat AddModuleScore function was used to assess Th17 skin scores (Supplementary Table 2) (11), PreP score (Supplementary Table 3) (9), and TCAT analysis (20) within IL23R enriched CD4 clusters.

### Serum analysis

Olink (Thermo Fisher Scientific, Waltham, Mass) proximity extension assay (Target-48 panel) was used to quantify plasma cytokines and chemokines as described previously (25). Analytes were measured from 1 μL of human plasma using an Olink Signature Q-100 instrument according to manufacturer’s protocol. Serum levels of IL-17A and IFN-y were analyzed using Spearman correlations and Wilcoxon matched pairs test.

## Data availability

Sequencing data is located at a link which will be added at time of publication. R codes for analysis will be made available upon request.

## Author contributions

B.N.W. and D.A.R conceptualized project. B.N.W. and D.A.R provided funding. K.E.M. and I.A. generated data. K.E.M. analyzed data. A.S. processed samples and recruited participants. L.P.C., A.P., L.B., J.F.M. and M.D.C. recruited participants and participated in data collection.

J.K. and M.G provided helpful advice. K.E.M., D.A.R., and B.N.W. drafted the manuscript, and all authors edited the manuscript.

## Acknowledgements

We thank the Brigham and Women’s Hospital Center for Cellular Profiling for cell sorting and scRNA sequencing preparation and assistance. Funding for this study was provided in part by NHLBI K23 HL159276-01 (to B.N.W.), AHA 21CDA851511 (to B.N.W.), NIAMS P30 AR070253, and Burroughs Wellcome Fund Career Award in Medical Sciences (to D.A.R.). This investigator-initiated study (MiNIMIA) was supported by an independent research grant from Sun Pharma.

---

**Supplemental Figure 1.**
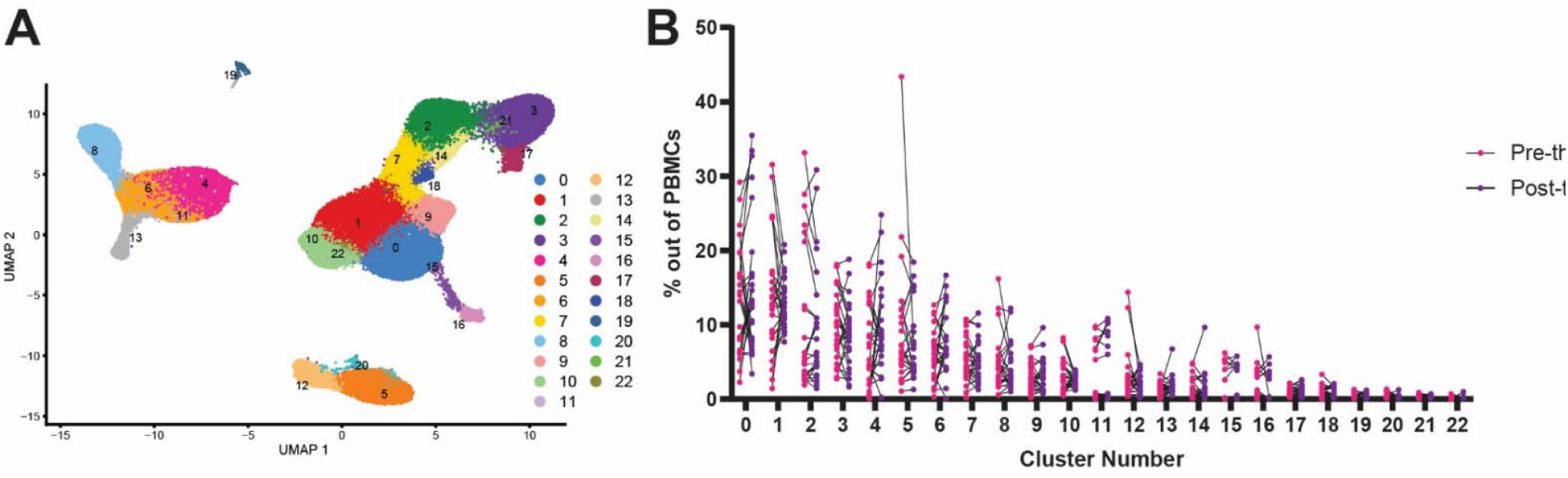
Longitudinal quantification of blood immune cell populations before and after IL-23 blockade. A) UMAP of scRNAseq of PBMCs from 27 psoriasis patients before and after IL-23 blockade therapy. B) Pre vs post proportions of PBMC clusters before vs after IL-23 blockade therapy. No changes are significant p > 0.05 by Wilcoxon test.

**Supplementary Figure 2.**
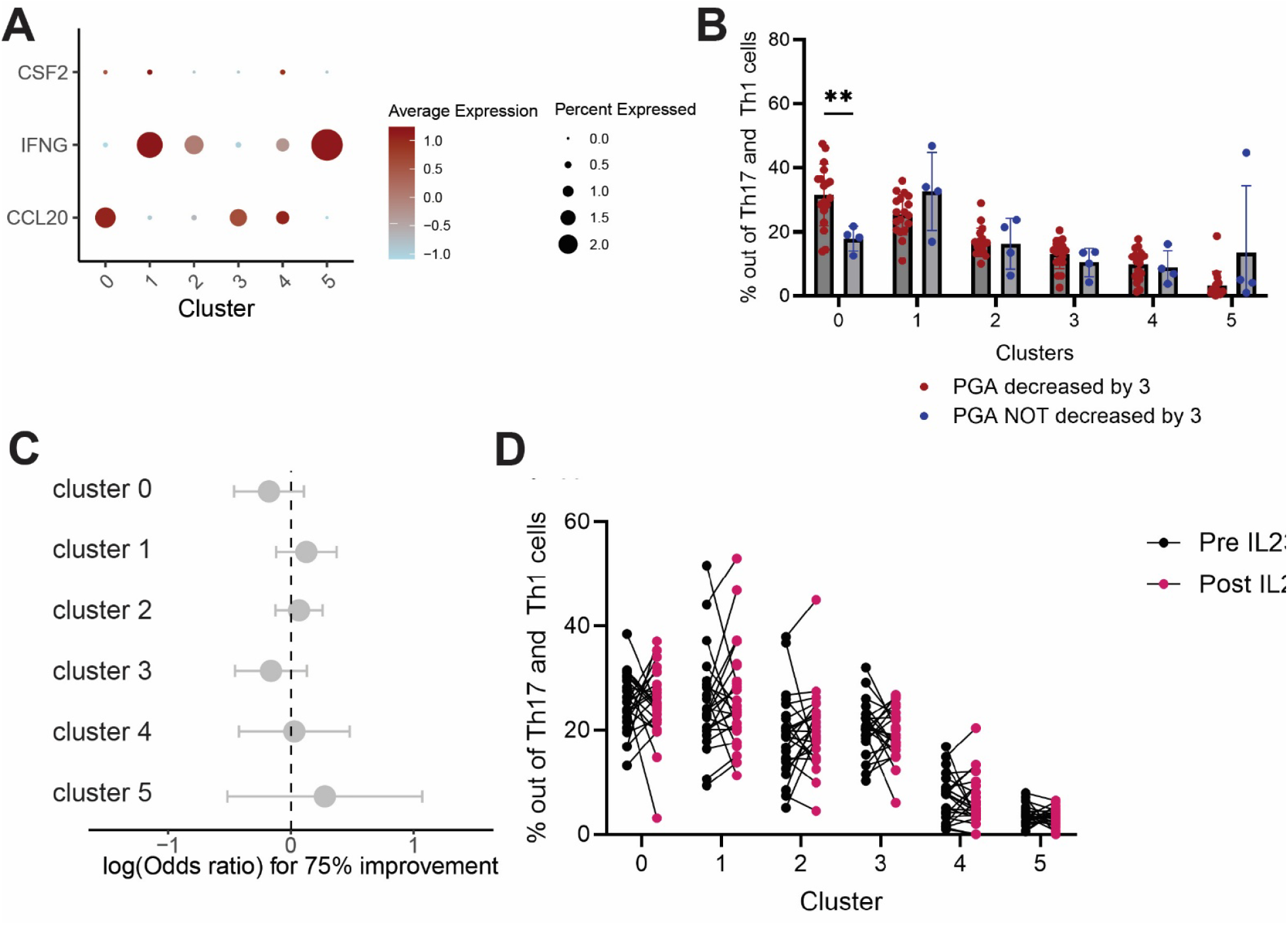
A) Gene expression within IL23R enriched CD4+ T cells. B) Proportion of each cluster divided by PASI 75. C) MASC analysis for association with PASI 75 among IL23R enriched cells after treatment. D) Pre vs post proportions of PBMC clusters before vs after IL-23 blockade therapy. All p values >0.05 by Wilcoxon test.

## References

1. Gelfand JM. Psoriasis — More Progress but More Questions. New England Journal of Medicine. 2024;390(6):561-2. doi: doi:10.1056/NEJMe2314345.

2. Kim J, Krueger JG. Highly Effective New Treatments for Psoriasis Target the IL-23/Type 17 T Cell Autoimmune Axis. Annu Rev Med. 2017;68:255–69. Epub 2016/10/01. doi: 10.1146/annurev-med-042915-103905. PubMed PMID: 27686018.

3. Gelfand JM, Neimann AL, Shin DB, Wang X, Margolis DJ, Troxel AB. Risk of myocardial infarction in patients with psoriasis. Jama. 2006;296(14):1735–41. Epub 2006/10/13. doi: 10.1001/jama.296.14.1735. PubMed PMID: 17032986.

4. Kim J, Krueger JG. The immunopathogenesis of psoriasis. Dermatol Clin. 2015;33(1):13–23. Epub 2014/11/22. doi: 10.1016/j.det.2014.09.002. PubMed PMID: 25412780.

5. Krueger JG, Ferris LK, Menter A, Wagner F, White A, Visvanathan S, Lalovic B, Aslanyan S, Wang EE, Hall D, Solinger A, Padula S, Scholl P. Anti-IL-23A mAb BI 655066 for treatment of moderate-to-severe psoriasis: Safety, efficacy, pharmacokinetics, and biomarker results of a single-rising-dose, randomized, double-blind, placebo-controlled trial. J Allergy Clin Immunol. 2015;136(1):116-24.e7. Epub 2015/03/15. doi: 10.1016/j.jaci.2015.01.018. PubMed PMID: 25769911.

6. Blaschitz C, Raffatellu M. Th17 Cytokines and the Gut Mucosal Barrier. Journal of Clinical Immunology. 2010;30(2):196–203. doi: 10.1007/s10875-010-9368-7.

7. Hernández-Santos N, Gaffen Sarah L. Th17 Cells in Immunity to *Candida albicans*. Cell Host & Microbe. 2012;11(5):425–35. doi: 10.1016/j.chom.2012.04.008.

8. Yen D, Cheung J, Scheerens H, Poulet F, McClanahan T, McKenzie B, Kleinschek MA, Owyang A, Mattson J, Blumenschein W, Murphy E, Sathe M, Cua DJ, Kastelein RA, Rennick D. IL-23 is essential for T cell–mediated colitis and promotes inflammation via IL-17 and IL-6. The Journal of Clinical Investigation. 2006;116(5):1310–6. doi: 10.1172/JCI21404.

9. Hu D, Notarbartolo S, Croonenborghs T, Patel B, Cialic R, Yang T-H, Aschenbrenner D, Andersson KM, Gattorno M, Pham M, Kivisakk P, Pierre IV, Lee Y, Kiani K, Bokarewa M, Tjon E, Pochet N, Sallusto F, Kuchroo VK, Weiner HL. Transcriptional signature of human pro-inflammatory TH17 cells identifies reduced IL10 gene expression in multiple sclerosis. Nature Communications. 2017;8(1):1600. doi: 10.1038/s41467-017-01571-8.

10. Zhao J, Lu Q, Liu Y, Shi Z, Hu L, Zeng Z, Tu Y, Xiao Z, Xu Q. Th17 Cells in Inflammatory Bowel Disease: Cytokines, Plasticity, and Therapies. J Immunol Res. 2021;2021:8816041. Epub 2021/02/09. doi: 10.1155/2021/8816041. PubMed PMID: 33553436; PMCID: PMC7846404 publication of this paper.

11. Kim J, Lee J, Kim HJ, Kameyama N, Nazarian R, Der E, Cohen S, Guttman-Yassky E, Putterman C, Krueger JG. Single-cell transcriptomics applied to emigrating cells from psoriasis elucidate pathogenic versus regulatory immune cell subsets. J Allergy Clin Immunol. 2021;148(5):1281–92. Epub 2021/05/02. doi: 10.1016/j.jaci.2021.04.021. PubMed PMID: 33932468; PMCID: PMC8553817.

12. Kim J, Lee J, Lee J, Kim K, Li X, Zhou W, Cao J, Krueger JG. Psoriasis harbors multiple pathogenic type 17 T-cell subsets: Selective modulation by risankizumab. J Allergy Clin Immunol. 2025;155(6):1898–912. Epub 2025/02/21. doi: 10.1016/j.jaci.2025.02.008. PubMed PMID: 39978685; PMCID: PMC12145251.

13. Langley RG, Ellis CN. Evaluating psoriasis with Psoriasis Area and Severity Index, Psoriasis Global Assessment, and Lattice System Physician’s Global Assessment. Journal of the American Academy of Dermatology. 2004;51(4):563–9. doi: 10.1016/j.jaad.2004.04.012.

14. Korsunsky I, Millard N, Fan J, Slowikowski K, Zhang F, Wei K, Baglaenko Y, Brenner M, Loh P-r, Raychaudhuri S. Fast, sensitive and accurate integration of single-cell data with Harmony. Nature Methods. 2019;16(12):1289–96. doi: 10.1038/s41592-019-0619-0.

15. Law C, Wacleche VS, Cao Y, Pillai A, Sowerby J, Hancock B, Horisberger A, Bracero S, Skidanova V, Li Z, Adejoorin I, Dillon E, Benque IJ, Nunez DP, Simmons DP, Keegan J, Chen L, Baker T, Brohawn PZ, Al-Mossawi H, Hao L-Y, Jones B, Rao N, Qu Y, Alves SE, Albrecht J, Anolik JH, Apruzzese W, Barnas JL, Bathon JM, Ben-Artzi A, Boyce BF, Boyle DL, Bridges SL, Bykerk VP, Campbell D, Ceponis A, Chicoine A, Curtis M, Deane KD, DiCarlo E, Donlin LT, Dunn P, Filer A, Carr H, Firestein GS, Forbess L, Geraldino-Pardilla L, Goodman SM, Gravallese EM, Gregersen PK, Guthridge JM, Gutierrez-Arcelus M, Holers VM, Horowitz D, Hughes LB, Ivashkiv LB, Ishigaki K, James JA, Jonsson AH, Kang JB, Keras G, Korsunsky I, Lakhanpal A, Lederer JA, Lewis MJ, Li Y, Liao K, Mandelin AM, Mantel I, Marks KE, Maybury M, McDavid A, McGeachy MJ, Mears JR, Meednu N, Millard N, Moreland L, Nayar S, Nerviani A, Orange DE, Perlman H, Pitzalis C, Rangel-Moreno J, Raychaudhuri S, Raza K, Reshef Y, Ritchlin C, Rivellese F, Robinson WH, Rumker L, Sahbudin I, Sakaue S, Seifert JA, Scheel-Toellner D, Singaraju A, Slowikowski K, Smith M, Tabechian D, Utz PJ, Watts GFM, Wei K, Weinand K, Weisenfeld D, Weisman M, Xiao Q, Zhang F, Zhu Z, Cordle A, Wyse A, Jonsson AH, Shaw KS, Vleugels RA, Massarotti E, Costenbader KH, Brenner MB, Lederer JA, Hultquist JF, Choi J, Rao DA, Accelerating Medicines Partnership RASLEN. Interferon subverts an AHR–JUN axis to promote CXCL13+ T cells in lupus. Nature. 2024;631(8022):857–66. doi: 10.1038/s41586-024-07627-2.

16. Stritesky GL, Yeh N, Kaplan MH. IL-23 Promotes Maintenance but Not Commitment to the Th17 Lineage1. The Journal of Immunology. 2008;181(9):5948–55. doi: 10.4049/jimmunol.181.9.5948.

17. Reshef YA, Rumker L, Kang JB, Nathan A, Korsunsky I, Asgari S, Murray MB, Moody DB, Raychaudhuri S. Co-varying neighborhood analysis identifies cell populations associated with phenotypes of interest from single-cell transcriptomics. Nature Biotechnology. 2022;40(3):355–63. doi: 10.1038/s41587-021-01066-4.

18. Carlin CS, Feldman SR, Krueger JG, Menter A, Krueger GG. A 50% reduction in the Psoriasis Area and Severity Index (PASI 50) is a clinically significant endpoint in the assessment of psoriasis. Journal of the American Academy of Dermatology. 2004;50(6):859–66. doi: 10.1016/j.jaad.2003.09.014.

19. Fonseka CY, Rao DA, Teslovich NC, Korsunsky I, Hannes SK, Slowikowski K, Gurish MF, Donlin LT, Lederer JA, Weinblatt ME, Massarotti EM, Coblyn JS, Helfgott SM, Todd DJ, Bykerk VP, Karlson EW, Ermann J, Lee YC, Brenner MB, Raychaudhuri S. Mixed-effects association of single cells identifies an expanded effector CD4(+) T cell subset in rheumatoid arthritis. Science translational medicine. 2018;10(463). Epub 2018/10/20. doi: 10.1126/scitranslmed.aaq0305. PubMed PMID: 30333237; PMCID: PMC6448773.

20. Kotliar D, Curtis M, Agnew R, Weinand K, Nathan A, Baglaenko Y, Zhao Y, Sabeti PC, Rao DA, Raychaudhuri S. Reproducible single cell annotation of programs underlying T-cell subsets, activation states, and functions. bioRxiv. 2024:2024.05.03.592310. doi: 10.1101/2024.05.03.592310.

21. Bunte K, Beikler T. Th17 Cells and the IL-23/IL-17 Axis in the Pathogenesis of Periodontitis and Immune-Mediated Inflammatory Diseases. Int J Mol Sci. 2019;20(14). Epub 2019/07/13. doi: 10.3390/ijms20143394. PubMed PMID: 31295952; PMCID: PMC6679067.

22. Reich K, Papp KA, Blauvelt A, Tyring SK, Sinclair R, Thaçi D, Nograles K, Mehta A, Cichanowitz N, Li Q, Liu K, La Rosa C, Green S, Kimball AB. Tildrakizumab versus placebo or etanercept for chronic plaque psoriasis (reSURFACE 1 and reSURFACE 2): results from two randomised controlled, phase 3 trials. The Lancet. 2017;390(10091):276–88. doi: 10.1016/S0140-6736(17)31279-5.

23. Heim J, Vasquez JG, Bhutani T, Koo J, Mathew J, Gogineni R, Ferro T, Bhatia N. Tildrakizumab Real-World Effectiveness and Safety Over 64 Weeks in Patients With Moderate-to-Severe Plaque Psoriasis. J Drugs Dermatol. 2024;23(8):612–8. Epub 2024/08/02. doi: 10.36849/jdd.8217. PubMed PMID: 39093661.

24. Hao Y, Hao S, Andersen-Nissen E, Mauck WM, Zheng S, Butler A, Lee MJ, Wilk AJ, Darby C, Zager M, Hoffman P, Stoeckius M, Papalexi E, Mimitou EP, Jain J, Srivastava A, Stuart T, Fleming LM, Yeung B, Rogers AJ, McElrath JM, Blish CA, Gottardo R, Smibert P, Satija R. Integrated analysis of multimodal single-cell data. Cell. 2021;184(13):3573-87.e29. doi: 10.1016/j.cell.2021.04.048.

25. Brodeur KE, Liu M, Ibanez D, de Groot MJ, Chen L, Du Y, Seyal E, Laza-Briviesca R, Baker A, Chang JC, Chang MH, Day-Lewis M, Dedeoglu F, Dionne A, de Ferranti SD, Friedman KG, Halyabar O, Lo MS, Meidan E, Sundel RP, Henderson LA, Nigrovic PA, Newburger JW, Son MB, Lee PY. Elevation of IL-17 Cytokines Distinguishes Kawasaki Disease From Other Pediatric Inflammatory Disorders. Arthritis & Rheumatology. 2024;76(2):285–92. doi: 10.1002/art.42680.

